# Combustible and electronic cigarette exposures increase ACE2 activity and SARS-CoV-2 Spike binding

**DOI:** 10.1101/2021.06.04.447156

**Authors:** Arunava Ghosh, Vishruth Girish, Monet Lou Yuan, Raymond D. Coakley, Neil E. Alexis, Erin L. Sausville, Anand Vasudevan, Alexander R. Chait, Jason M. Sheltzer, Robert Tarran

## Abstract

The outbreak of coronavirus disease 2019 (COVID-19) has extensively impacted global health. The causative pathogen, severe acute respiratory syndrome coronavirus 2 (SARS-CoV-2), binds to the angiotensin-converting enzyme 2 (ACE2) receptor, a transmembrane metallo-carboxypeptidase that is expressed in both membrane-anchored (mACE2) and soluble (sACE2) forms in the lung. Tobacco use has been speculated as a vulnerability factor for contracting SARS-CoV-2 infection and subsequent disease severity, whilst electronic cigarettes (e-cigarettes) have been shown to induce harmful proteomic and immune changes in the lungs of vapers. We therefore tested the hypothesis that combustible tobacco (e.g. cigarettes) and non-combustible e-cigarettes could affect ACE2 activity and subsequent SARS-CoV-2 infection. We observed that sACE2 activity was significantly higher in bronchoalveolar lavage fluid from both smokers and vapers compared to age-matched non-smokers. Exposure to cigarette smoke increased ACE2 levels, mACE2 activity, and sACE2 in primary bronchial epithelial cultures. Finally, treatment with either cigarette smoke condensate or JUUL e-liquid increased infections with a spike-coated SARS-CoV-2 pseudovirus. Overall, these observations suggest that tobacco product use elevates ACE2 activity and increases the potential for SARS-CoV-2 infection through enhanced spike protein binding.

## To the Editor

The outbreak of coronavirus disease 2019 (COVID-19) has extensively impacted global health. The causative pathogen, severe acute respiratory syndrome coronavirus 2 (SARS-CoV-2), binds to the angiotensin-converting enzyme 2 (ACE2) receptor, a transmembrane metallo-carboxypeptidase that is expressed in both membrane-anchored (mACE2) and soluble (sACE2) forms in the lung. Although mACE2 is responsible for viral entry, recent observations also suggest a role of sACE2 in infection (1). Tobacco use has been speculated as a vulnerability factor for contracting SARS-CoV-2 infection and subsequent disease severity (2, 3), whilst electronic cigarettes (e-cigarettes) have been shown to induce harmful proteomic and immune changes in the lungs of vapers (Ghosh A et al., AJRCCM, 2018). We therefore tested the hypothesis that combustible tobacco (e.g. cigarettes) and non-combustible e-cigarettes could affect ACE2 activity and subsequent SARS-CoV-2 infection.

We first evaluated sACE2 activity in concentrated bronchoalveolar lavage fluid (BALF) samples from non-smokers (age, 28.5 ± 9.5 years; body mass index [BMI], 28.8 ± 7.2 kg/m^2^; percentage predicted FVC [ppFVC], 98.0 ± 25.0; ppFEV1, 101.6 ± 14.5); smokers (age, 34.4 ± 8.2 years; BMI, 27.5 ± 7.8 kg/m^2^; ppFVC, 104.2 ± 11.2; ppFEV1, 97.9 ± 16.1) and vapers (age, 29.3 ± 9.1 years; BMI, 29.6 ± 8.0 kg/m^2^; ppFVC, 106.2 ± 11.2; ppFEV1, 104.5 ± 7.7). We measured cleavage of an ACE2-specific fluorogenic peptide (Vivitide, SFQ-3819-PI) by microplate reader to evaluate ACE2 activity as described (4). sACE2 activity was significantly increased in BALF from both smokers and vapers compared to age-matched non-smokers (Figure 1A), indicating heightened vulnerability towards SARS-CoV-2 infection in both groups.

**Figure 1.**
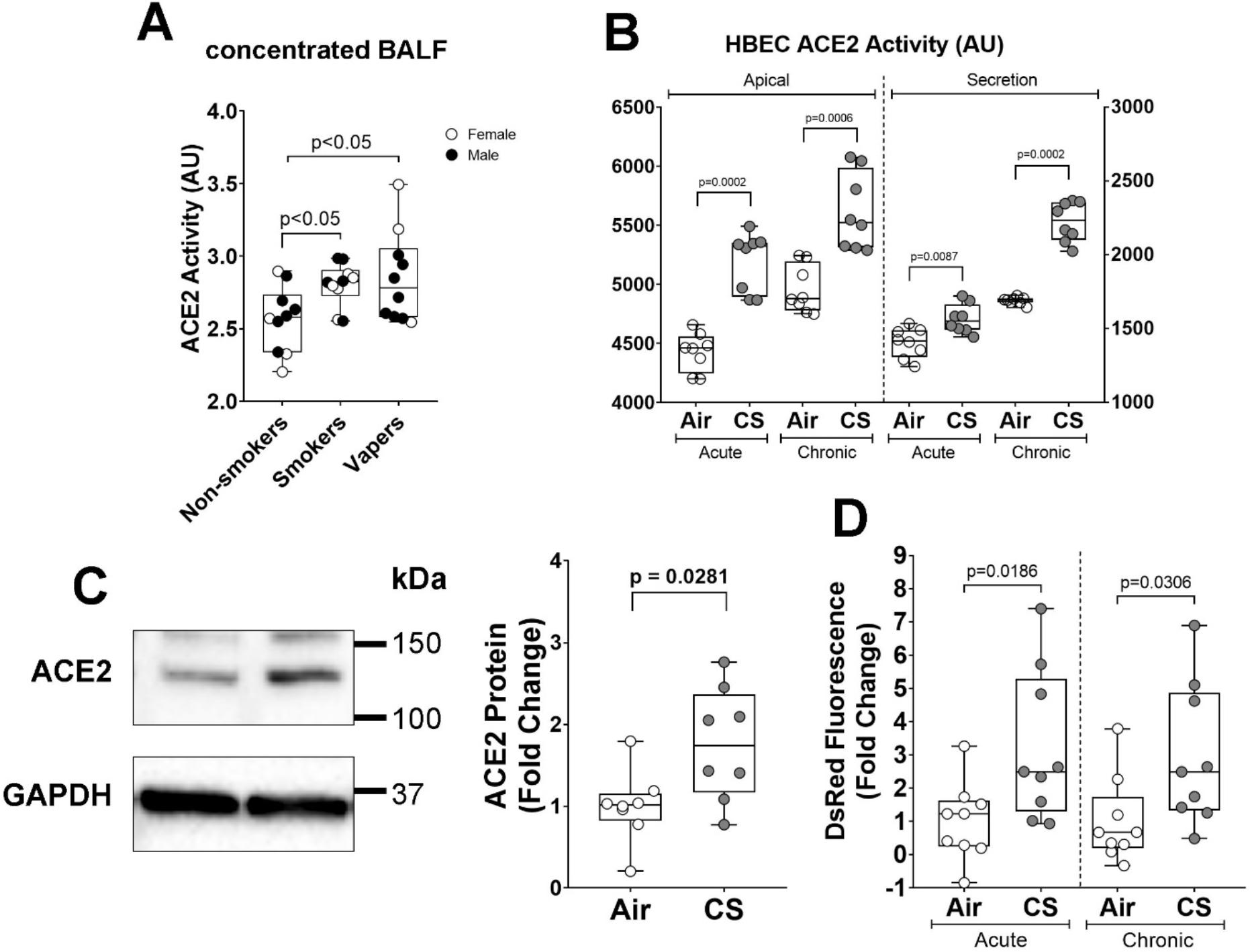
Tobacco smoking increases ACE2 activity and SARS-CoV-2 infection. Bronchoalveolar lavage fluids from non-smokers, smokers, and vapers (n=10 each) were concentrated and ACE2 activity was measured using 50 μM specific fluorogenic substrates. Cleaved substrate derived fluorescence as arbitrary unit (AU) following one hour of reaction was reported (A). Human bronchial epithelial cells (HBECs) were cultures at air-liquid interface (ALI) and exposed to smoke from one cigarette per day for one (acute) or four days (chronic). Apical surface was washed with PBS to collect the mucosal secretion and 50 μM fluorogenic substrate was added on the apical surface to evaluate apical ACE2 activity. ACE2 activity on apical side of acute and chronic smoke exposed cultures and corresponding apical washes (B) were reported. Activities are presented as accumulated fluorescence in arbitrary units (AU) of the cleaved products following 10 minutes of reaction. Whole cell lysates were collected from chronically cigarette smoke exposed cultures, and ACE2 and GAPDH protein expressions were evaluated by immunoblotting. Representative blots were shown and GAPDH normalized ACE2 expression in air and smoke exposed cultures as fold change (C) was reported. (D) Acute or chronically smoke exposed cultures were apically exposed to spike pseudovirus suspension and the resulting DsRed marker fluorescence was recorded after three days. Data represented as median, lower and upper quartiles; n=8-9 per group; p-values were provided on the graph.

To evaluate the impact of tobacco smoke on ACE2 activity, we grew primary human bronchial epithelial cultures (HBECs) at air-liquid interface (ALI) and exposed them to 14 puffs of smoke per day from Kentucky research cigarettes for one day (acute) and four days (chronic). Acute smoke exposure significantly increased both mACE2 and sACE2 activity (Figure 1B). Following chronic smoke exposure, both mACE2 and sACE2 activity remained significantly elevated compared to air controls (Figure 1B). Consistent with these observations, ACE2 protein was significantly increased in chronically smoke-exposed cultures compared to air exposed controls (Figure 1C), indicating that more receptors were potentially available for SARS-CoV-2 binding. As an orthogonal approach, we developed a pseudovirus using recombinant vesicular stomatitis virus ΔG (rVSV-ΔG, a gift from Pengfei Wang, Columbia University) that co-expressed the SARS-CoV-2 spike protein (Addgene, 145032 or 164583) and DsRed. We exposed HBECs to acute and chronic tobacco smoke, followed by infection with 3×10^4^ U/mL of pseudovirus, and measured DsRed expression by microplate reader. Cigarette smoke exposure significantly increased DsRed fluorescence (increased viral infection; Figure 1D). We then incubated chronically air- and smoke-HBECs with recombinant SARS-CoV-2 S1 protein. Consistent with the observed increase in ACE2 activity, smoke exposure caused a significant (20%, n=9, p=0.024) increase in S1 binding. Taken together, these data indicate that cigarette smoke exposure upregulates ACE2 activity and increases SARS-CoV-2 binding.

Next, we explored the impact of cigarette smoke condensate (CSC) and e-liquid exposure on pseudovirus infection in primary tracheobronchial (ATCC, PCS-300-010) and small airway epithelia (ATCC, PCS-301-010). Cells were cultured to confluence on plastic, treated with a range of different doses of CSC (Murty Pharmaceuticals) or JUUL Virginia Tobacco-flavored e-liquid (5% nicotine benzoate) for 24 hours, and infected with pseudovirus (2.5-5×10^5^ U/mL). We then used fluorescent microscopy and flow cytometry to visualize and quantify this infection respectively. Acute exposure to both CSC and JUUL e-liquid significantly increased DsRed fluorescence, indicating increased pseudovirus infection compared to control cells (Figure 2A-D). These data correlate with our BALF data (Figure 1A) and indicate that both cigarettes and e-cigarettes increase the potential for SARS-CoV-2 infection.

**Figure 2.**
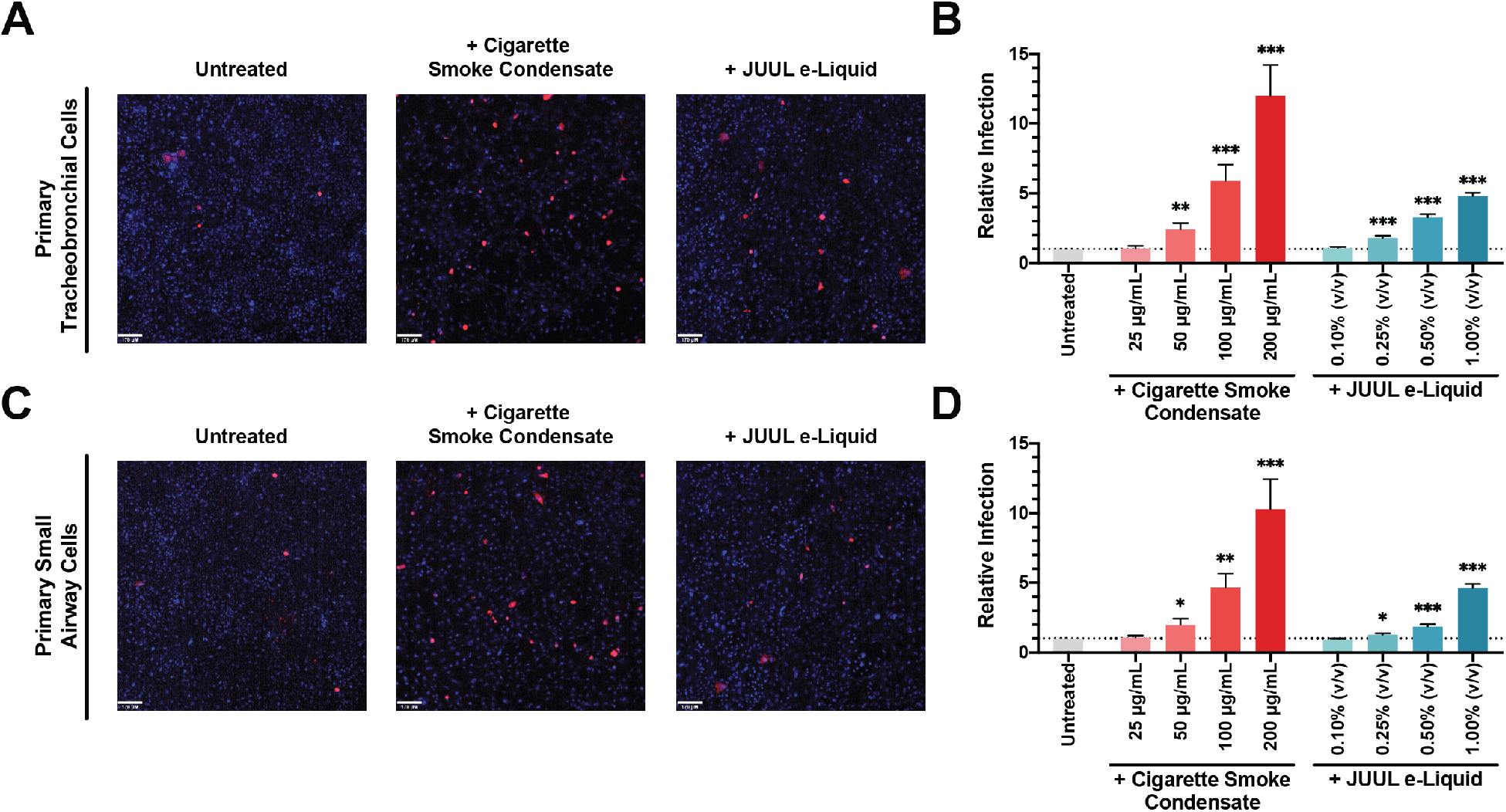
Cigarette Smoke Condensate Exposure Increases SARS-CoV-2 Pseudovirus Infection. Primary tracheobronchial (A) and small airway cells (C) were pre-treated with 200 μg/mL CSC (middle) and 1.00% JUUL e-Liquid (right) for 24 hours and challenged with SARS-CoV-2 pseudovirus. Following incubation for 48 hours, live cells were imaged on a spinning-disc confocal microscopy system (UltraVIEW Vox; PerkinElmer). Cell nuclei were stained with Hoechst dye and infected cells were shown in red. Flow cytometry quantification of relative SARS-CoV-2 pseudovirus infection in different doses of CSC and JUUL e-Liquid pre-treated primary tracheobronchial cells (B) and small airway cells (D) were reported as fold change; n = 11-12 trials. Bars represent mean ± SEM. Unpaired t test * p < 0.05; ** p < 0.005; *** p < 0.0005. Scale bars, 170 μm.

Several meta-analyses have been performed to look at tobacco use and COVID-19 disease severity, which have reported contradictory findings. A “nicotinic hypothesis” was proposed, which suggested that nicotine exerted a protective role against SARS-CoV-2 infection (5). Conversely, other reports confirmed a strong association between smoking and COVID-19 disease symptoms (2, 3). Increased ACE2 expression in smokers has been proposed as a risk factor for COVID-19 (6). Moreover, tobacco smoking is a risk factor for several other respiratory viruses including influenza (7). Our current study provides further *in vivo* evidence that sACE2 is upregulated in smokers’ and vapers’ BALF. However, our study has some limitations: First, our study was cross-sectional and of relatively low sample size, so we were unable to probe our data sets to understand the impact of gender, race and other factors. A second limitation of our study is that we only measured sACE2 *in vivo* and not mACE2. However, sACE2 facilitates SARS-CoV-2 infection through interactions with the renin-angiotensin system and elevated sACE2 in smokers and vapers is expected to contribute towards SARS-CoV-2 infection (1). The increase in sACE2 most likely reflects an overall increase in cellular ACE2 expression. Indeed, ACE2 mRNA expression is elevated in smokers’ pulmonary cells relative to those derived from non-smokers, suggesting that both membrane-associated and soluble ACE2 may be increased (8). Moreover, elevated sACE2 was positively associated with COVID-19 disease severity (9), demonstrating the utility of sACE2 as a marker of COVID-19 risk.

Nicotine, which is present in both combustible and electronic cigarettes, increases cellular Ca^2+^ levels (4). Elevated Ca^2+^ signaling is well-recognized to play a vital role in facilitating viral entry, gene and protein processing, and subsequent viral release (10). Our studies demonstrate that tobacco and e-liquid exposures increase both ACE2 and SARS-CoV-2 spike protein binding. Thus, we hypothesize that nicotine may be responsible for the increased spike protein binding and subsequent pseudovirus infection. However, the underlying mechanisms, and the purported link to nicotine will require additional study. We further speculate that acute but persistent increases in ACE2 and spike protein binding place tobacco users at risk of SARS-CoV-2 infection. Overall, these observations indicate that tobacco product use elevates ACE2 activity and increases the potential for SARS-CoV-2 infection through enhanced spike protein binding. Importantly, our results strongly urge consideration of vaping as a risk factor for COVID-19.

